# Negative frequency-dependent selection is frequently confounding

**DOI:** 10.1101/113324

**Authors:** Dustin Brisson

## Abstract

*This preprint has been reviewed and recommended by Peer Community in Evolutionary Biology (http://dx.doi.org/10.24072/pci.evolbiol.100024).* The existence of persistent genetic variation within natural populations presents an evolutionary problem as natural selection and genetic drift tend to erode genetic diversity. Models of balancing selection were developed to account for the high and sometimes extreme levels of polymorphism found in many natural populations. Negative frequency-dependent selection may be the most powerful selective force maintaining balanced natural polymorphisms but it is also commonly misinterpreted. The aim of this review is to clarify the processes underlying negative frequency-dependent selection, describe classes of natural polymorphisms that can and cannot result from these processes, and discuss observational and experimental data that can aid in accurately identifying the processes that generated or are maintain diversity in nature. Finally, I consider the importance of accurately describing the processes affecting genetic diversity within populations as it relates to research progress.

## Introduction

Natural diversity - the “endless forms most beautiful and most wonderful” [1] - is an enduring focus of both evolutionary biologists and nature lovers. The evolutionary processes that have generated or are maintaining many examples of diversity in nature, however, remain obscure and can be controversial [2]. The processes that result in persistent polymorphisms within populations demand a special explanation as both directional natural selection and genetic drift should eliminate alleles and thus erode genetic diversity [3–5]. Nevertheless, many examples of persistent polymorphisms occur in nature [6–11]. Models of balancing selection - including negative frequency-dependent selection, spatial or temporal habitat heterogeneity, and heterozygote advantage - provide theoretical frameworks of the processes that can account for persistent polymorphisms within populations. A core tenet of each balancing selection model is that the selective value of an allele – whether it is beneficial or detrimental – is dependent on the environmental context [12,13]. That is, alleles are advantageous and deleterious in different ecological contexts.

Negative frequency-dependent selection has been called the most powerful selective force maintaining balanced polymorphisms [14–17], with some proposing that a large proportion of natural genetic polymorphisms are maintained by selection favoring rare alleles [18]. Negative frequency-dependent selection occurs when the selective value of a variant relative to other variants is a function of its abundance in the population relative to other variants such that its relative fitness increases as the relative abundance, or frequency, of the variant decreases [19](please see [20,21] for foundational mathematical descriptions and assumptions of this process). That is, rare variants have a selective advantage specifically because of their rarity while common variants are disadvantaged because of their commonness. Thus, negative frequency-dependent selection has the potential to maintain polymorphisms within populations because relatively rare variants have a selective advantage over more common variants and thus tend to increase in frequency and avoid local extinction. Negative frequency-dependent selection models are a narrow subset of a broad field of models describing the impact of the frequency of variants on natural selection; the overwhelming majority of this broad field is beyond the scope of the concepts addressed here. Here, I focus on natural polymorphisms that can be explained by negative frequency-dependent selection, where genetic diversity is maintained when a variant becomes disadvantageous as it becomes more frequent, and polymorphisms that are more accurately explained by other process.

Numerous ecological interactions can result in a selective advantage for relatively rare alleles including sexual selection, parasite or predator preferences, and resource competition. In fact, each of these mechanisms has been shown to create a selective advantage for rare alleles that has resulted in persistent polymorphisms in multiple natural populations [10,19,22–24]. While ecological context and natural history determine the proximate ecological mechanism affecting the differential survival or reproduction of variants in a population, changes in relative survival or reproduction must be negatively correlated with variant frequency for negative frequency-dependent selection to maintain natural polymorphisms. In a classic example, color polymorphisms are maintained in natural populations of *Cepaea nemoralis* snails by negative frequency-dependent selection because their predators, the song thrush (*Turdus philomelos*), form a search image for the most common morph causing a much greater predation pressure on the common than the rare morph [23,25]. The rare morph can increase in frequency due to the relaxed predation pressure until it becomes common, resulting in a search image switch that now targets the new common morph, a process that maintains this polymorphism in *C. nemoralis* populations. Two luminaries in population genetics - R. Fisher and S. Wright - have also demonstrated the power of negative frequency-dependent selection to maintain diversity in natural systems. Wright famously demonstrated that self-incompatibility alleles, a genetic mechanism in plants to prevent inbreeding, are incredibly diverse because pollen containing a rare allele is more likely to find a receptive mate than pollen containing a common allele [26–28]. Thus, plants with rare alleles have a selective advantage (**Figure 1**). Similarly, Fisher’s principle demonstrates that males and females are equally frequent because, if one sex were more frequent, the alternate sex would enjoy a *per capita* reproductive advantage [22,29].

**Figure 1.**
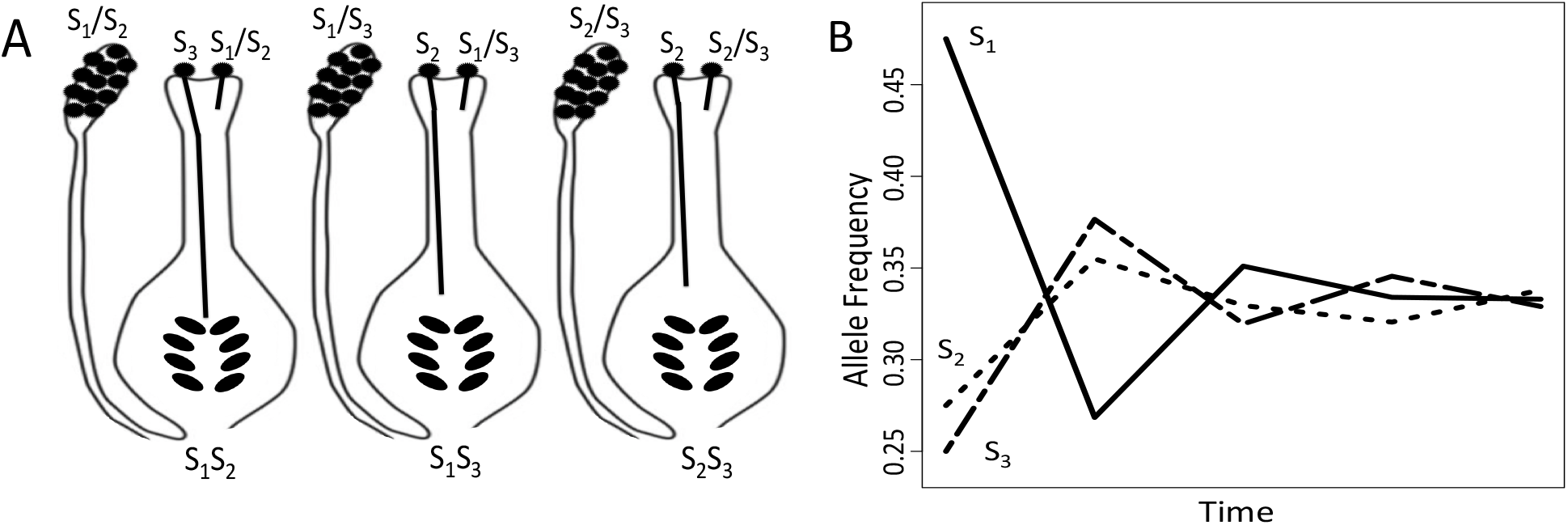
**A**. A cartoon depiction of the self-incompatibility allele model modified from [19]. For simplicity, the plant population represented has only three alleles although most populations maintain 10s or 100s of *S*-alleles. As describe in the classical model [19], *S_1_S_1_* homozygote plants produce only pollen containing the *S_1_* allele and can be pollinated only by pollen with the *S_2_* or *S*_*3*_ allele. Similarly, *S_2_S_2_* homozygotes produce *S_2_*-containing pollen and can be pollinated only by *S_1_*- or *S_3_*-containing pollen. Heterozygote plants can produce pollen with either of its two alleles but cannot be pollinated by either pollen variant. Alleles that are relatively rare in the population have a selective advantage over common alleles as pollen containing a rare allele is much more likely to pollinate a receptive ovule. In contrast, pollen containing a common allele is likely to attempt to pollinate a plant containing a common allele and be rejected, resulting in limited breeding success. **B**. The temporal dynamics of the alleles in this system are likely to fluctuate as expected when rare alleles have a selective advantage. For example, if 81% of the plants are homozygous *S_1_S_1_* (time 0), the *S_2_*-containing pollen (~10%) has a high probability of finding an *S_1_S_1_* plant and successfully breeding. By contrast, the *S*_*x*_-containing pollen is highly unlikely to find a *S_2_S_2_* homozygote (~1% of all plants), resulting in very low breeding success. Due to the limited breeding success, the *S*_*1*_ allele will decrease in frequency until *S_1_*-containing pollen is rare and becomes more likely to find a receptive mate. These dynamics occur because a pollen grain with a common allele will be limited in terms of mates, while a pollen grain with a rare allele will not. Hence, plants with rare alleles have a selective advantage in terms of mating (see formal model in [19]).

The many incontrovertible demonstrations of the power of negative frequency-dependent selection to maintain polymorphisms in nature have led some to suggest that it is a “pervasive” force maintaining natural diversity [21]. The pervasiveness of negative frequency-dependent selection is further supported by the perception that “nearly every [selective agent] works in a way liable to produce frequency-dependent selection of the kind that favours rare phenotypes and hinders common ones” [21]. Although negative frequency-dependent selection may be a “powerful, perhaps a dominant, factor maintaining genetic diversity” within populations [21], many natural polymorphisms are maintained by other evolutionary processes [30–35]. Still, many natural polymorphisms have been assumed to result from negative frequency-dependent selection even when the theoretical framework and data from the system are inconsistent with the processes of selection favoring relatively rare variants. In this essay, I describe several patterns of allele dynamics that are commonly described in the literature as resulting from negative frequency-dependent selection despite data demonstrating that other explanatory processes are causative. These processes include allelic diversity resulting from directional selection within a changing ecological context, density-dependent population regulation, other models of balancing selection, and aspects of community ecology. I will discuss concepts and experiments that can aid in identifying the processes underlying patterns of allele dynamics and suggest that accurately identifying the evolutionary process underlying natural patterns facilitates the development of hypotheses and future experiments to determine the ecological interactions or molecular mechanisms at the root of the process.

### Directional selection attributed to negative frequency-dependent selection

As a broad concept, negative frequency-dependent selection may be the “most intuitively obvious explanation” of polymorphisms in nature [36]. However, the original concept becomes ambiguous, complex, and even controversial as a result of differing applications in both theoretical and empirical work [37]. Even some of the greatest thinkers in evolutionary biology have explained scenarios where the selective values of alleles are independent of their relative abundance through a negative frequency-dependent selection framework. A prominent example comes from a sweeping and influential essay by JBS Haldane where he suggested several “lines of thought” concerning infectious diseases as a major selective force in metazoan evolution [38]. Contrary to his assertion that “many or all” of these ideas “may prove to be sterile,” most have been “followed profitably” (very profitably indeed). However, the negative frequency-dependent selection framework described in this essay appears to be one of the few unsound lines of thought. In this framework, Haldane suggested that a host with a rare defensive phenotype has a selective advantage in the face of highly-adapted pathogens, “For just because of its rarity it will be resistant to diseases which attack the majority of its fellows.” That is, the adapted pathogen has evolved mechanisms to overcome the common defensive phenotypes in host populations but cannot overcome rare defensive phenotypes. Thus, hosts expressing rare but effective defensive phenotypes enjoy a selective advantage over hosts expressing common but exploitable defenses.

The scenario described by Haldane, however, confounds natural selection favoring a specific (*effective*) phenotype in the current environment with a selective advantage resulting from rarity. Haldane’s escape variants have a selective advantage because they cannot be subverted by the pathogen, not because they are rare. Further, the novel defensive phenotype has not yet lost its efficacy against the pathogen not because it is rare, but because it is novel. This point can be illustrated by extending this line of thought to allow migration of many individuals expressing a novel and effective defensive phenotype. These migrants would enjoy the same selective advantage over the previously common resident phenotype, regardless of frequency of the novel phenotype in the population immediately following the mass-migration event. The evolutionary dynamics occurring in this framework do not occur because of rare advantage and, in most cases, will not result in a balanced polymorphism. These evolutionary dynamics are more likely the result of directional selection in a continuously changing environment [39–43]. These two processes - negative frequency-dependent selection and selection in a changing environment - can potentially be distinguished by artificially manipulating variant frequencies or by introducing a previously common but now extinct variant into a controlled population.

The genetic diversity of haemagglutinin (HA) glycoproteins in the influenza virus is another conspicuous example of selection in a changing environment often confounded with negative frequency-dependent selection. The dynamics of HA alleles change over time such that rare alleles enter the population, rise to high population sizes, and subsequently decline toward extinction [44–46]. The strains expressing a numerically common allele have relatively low fitness and decline in frequency because there are few hosts still susceptible to this strain as hosts acquire immunity to strains with which they have been previously infected [47–49]. By contrast, strains expressing numerically rare alleles have many susceptible hosts available and enjoy high rates of secondary infections per infected host causing an increase in frequency [48,49]. While there is undoubtedly strong selection at the HA locus, the selective advantage is derived not from relative rarity but from antigenic novelty [48,50–52], similar to Haldane’s example. The presence or frequency of alternative HA alleles does not affect the fitness (growth rate) or temporal dynamics of the alleles. That is, the population dynamics of a numerically rare allele is the same if the host population is already plagued by other numerically common strains (0.0001% when one novel allele enters a population of 10^6^ infected hosts) and if it enters a host population in which no other influenza strain is circulating (100% when one novel allele enters a previous uninfected host population) (**Figure 2**). As the selective value of the allele is conditioned on the absolute abundance - but not the relative abundance - of the allele, it is unlikely that negative frequency-dependent selection is the evolutionary process underlying the polymorphism commonly observed at the HA locus. More likely, the common variant is changing its own environment such that there are few susceptible hosts in which new infections can establish, but is not affecting the environment of alternative variants.

**Figure 2.**
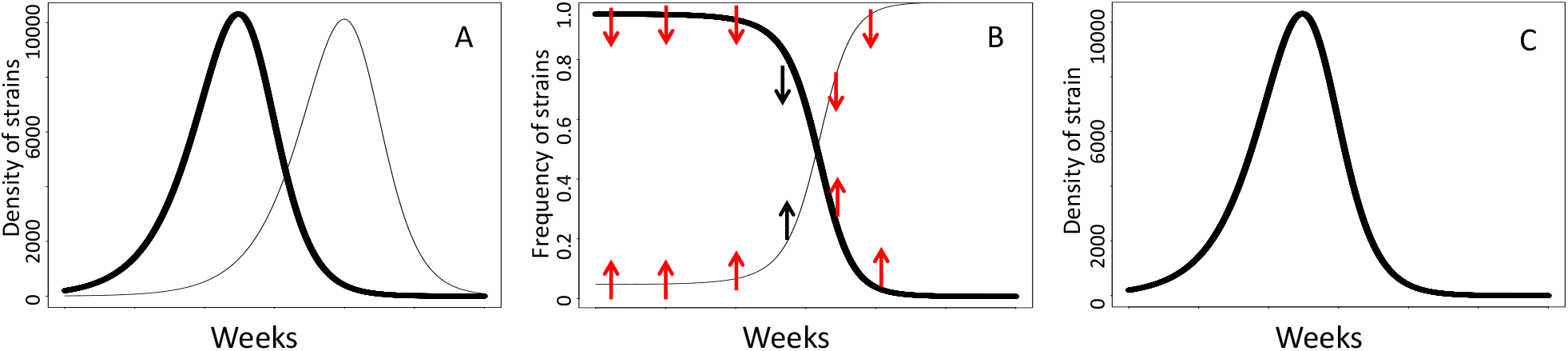
Influenza virus carrying rare HA or NA alleles do not have a selective advantage because they are relatively rare – a necessary condition of negative frequency-dependent selection – but because they are numerically rare compared to the number of susceptible hosts. **A**. The population dynamics of two influenza strains (dark and light lines) that enter a host population sequentially. Both strains increase numerically when they are numerically rare, but not relatively rare, and decrease after they become numerically common. For example, the maximal rate of increase of the first strain occurs prior to the second strain entering the population, despite remaining at a maximal relative abundance (100%). **B**. The relative frequencies of the two influenza strains through time. If negative frequency-dependent selection were affecting the relative abundances of these strains, the common strain at time=0 (dark line) should have lower fitness than the rare strain (light line). However, the *per capita* rate of increase of the common strain remains high until it has substantially reduced the number of susceptible hosts, regardless of its relative abundance. The arrows indicate expected effect of negative frequency-dependent selection on the relative fitness of each strain given its relative abundance. Red arrows indicate the time periods when the expectations of negative frequency-dependent selection are not satisfied; black arrows indicate time periods when negative frequency-dependent selection expectations are satisfied. **C**. The *per capita* rate of increase (numerical growth rate) and the population dynamics of each strain have the same temporal patterns in the absence of the alternative strain. Strain 1 remains at 100% frequencies throughout the time period, suggesting that relative abundance does not drive of changes in relative fitness.

### Density-dependent fitness dynamics attributed to negative frequency-dependent selection

A preeminent evolutionary biologist, R.C. Lewontin, suggested that negative frequency-dependent selection should be pervasive because, whenever “a genotype is its own worst enemy, its fitness will decrease as it becomes more common” [3]. As similar variants occupy similar niches and are commonly their own worst enemy, this logic suggests that negative frequency-dependent selection should indeed be pervasive. However, “common” in this case refers not to relative abundance but absolute abundance. For example, the fitness (growth rate) of individuals within a monomorphic population, one in which the frequency of a genotype is always at 100%, decreases as it “becomes more common” in absolute abundance. Further, relatively rare variants suffer negative fitness effects in proportion to the absolute abundance of their numerically common competitors such that relative rarity may not provide a selective advantage.

There is an extensive literature describing fitness (growth rate) as a function of the absolute abundance of each variant in a population [53–57]. The above scenario can be characterized using classical Logistic growth models that include competition among variants such that “a genotype is its own worst enemy” (Lotka-Volterra models) (**eq. 1**). The growth rates of the variants in these models are a function of the absolute abundance of each variant – discounted by their competitive abilities (*α_ij_*) – with respect to the carrying capacity (*K*), but are not conditioned on the abundance of the variants relative to each other. An interesting body of literature uses this modeling framework to describe the generation and maintenance of polymorphisms not through negative frequency-dependent selection mechanisms but through disruptive selection conditioned on the strength of competitive interactions and the abundance of each variant (ex [58]).

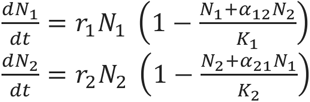

It is often challenging to distinguish the effect of numerical rarity from relatively rarity on the selective value of an allele through observations of patterns of allelic diversity. Experimental manipulations of the carrying capacity (*K*), potentially through resource supplementation, can assuage the reductions in relative fitness experienced by common variants that result from high densities without altering relative frequencies. In these experiments, the relative fitness of common variants should increase if the effects are associated with density while the relative fitness of the common and rare variants should not be altered if the allelic diversity is maintained by negative frequency-dependent selection.

### Multiple niche polymorphisms attributed to negative frequency-dependent selection

In the multiple niche selection model of balancing selection, the selective value of an allele is conditioned on their ability to exploit different environmental features in a heterogeneous habitat [59,60]. Multi-niche selection maintains multiple variants in a population if each variant has a selective advantage in some available habitats while other variants are superior in other habitats. This idea – that environmentally variable selection can result in balanced polymorphisms – has a long history in the literature in which the foundational idea is stated by Dobzhansky [12]. Although incontrovertible examples of multiniche selection maintaining polymorphism in natural populations are relatively rare, correct inference of the process resulting in balancing selection is necessary to generate hypotheses and controlled experiments to determine the underlying ecological interactions or molecular mechanisms causing the process.

The study of pattern, in isolation from the evolutionary processes that generated it, is not likely to advance general theories nor an understanding of a specific system [61]. However, determining the processes responsible for balanced polymorphism patterns observed in nature is a difficult task [17,62–64]. The balanced polymorphism at the outer surface protein C (*ospC*) locus in populations of *Borrelia burgdorferi*, the cause of human Lyme disease, provides a fitting example. Although the function of OspC remains unclear [65–70], the within-population diversity at this locus bears all the hallmarks of balancing selection - large numbers of alleles in all local populations; allele frequencies that are more even than expected for neutrally evolving loci; and genetic evidence of an ancient polymorphism [34,71–75].

Negative frequency-dependent selection and multi-niche selection have both been proposed as processes maintaining the *ospC* polymorphisms, and both frameworks have empirical support [71,73,76–78]. The negative frequency-dependent selection model suggests that the polymorphism can be maintained if previously infected hosts are immune to subsequent infections by the same OspC variant but susceptible to novel variants, a molecular mechanism that has been demonstrated in laboratory animals [79,80, but see, 81]. However, in this scenario the frequency or even presence of alternative OspC variants does not affect the number of susceptible hosts for the common strain, similar to the influenza example, arguing against negative frequency-dependent selection as an evolutionary process maintaining *ospC* polymorphisms. Further, negative frequency-dependent selection is most effective when few hosts remain susceptible to the common *ospC* types, a pattern that is not observed in natural data sets [34,82–85]. Studies investigating allelic diversity at *ospC* from natural hosts consistently demonstrate that most natural reservoir hosts, those that are regularly infected with *B. burgdorferi*, are rarely infected with all of the common *ospC* alleles [34,82,84,86]. Most hosts are, however, infected with a subset of the *ospC* alleles, as expected if each host species represented a different ecological niche [34,82,84,86]. Further, host individuals of the same species, including humans, are infected by the same subset of *ospC* alleles across both time and space [34,82,84,86–89]. The collective evidence suggests that the balanced *ospC* polymorphisms are more likely maintained by multi-niche selection - with each host species representing multiple niches [90], one for each *ospC* variant by which it can be infected - than by negative frequency-dependent selection. These results suggest that the mechanisms causing the balanced polymorphism are more likely to involve genotype-by-host species interactions than to involve a memory immune response mechanism that is conserved across vertebrate species.

It has been argued that “Selection in multiple niches is not an alternative to [negative] frequency-dependent selection…but a way of generating it” [21]. However, scenarios in which balanced polymorphisms can be maintained without a selective advantage favoring relatively rare variants are not uncommon, suggesting that these are two distinct evolutionary processes in at least some cases. To illustrate this point, image two variants occupy a heterogeneous habitat where each variant has a selective advantage in one niche but is disadvantaged in another, a classical multi-niche selection model [59,60]. Here we assume that the carrying capacity in niche A is much lower than the carrying capacity in niche B (K_A_=10; K_B_=10^5^). In this scenario, variant B - which has a competitive advantage in niche B - can retain a fitness advantage (a greater *per capita* growth rate) even when it is more common than variant A - which has a competitive advantage in niche A. For example, in a population with 90 variant B individuals and 10 variant A individuals, variant B has a rapid *per capita* rate of increase while variant A does not increase (**Figure 3**). Here, the relatively common variant B has a “selective advantage” over the relatively rare variant A due to multi-niche selection, which is independent of negative frequency-dependent selection. Depending on the parameter values in this model, a balanced polymorphism can be maintained in the absence of rare advantage.

**Figure 3.**
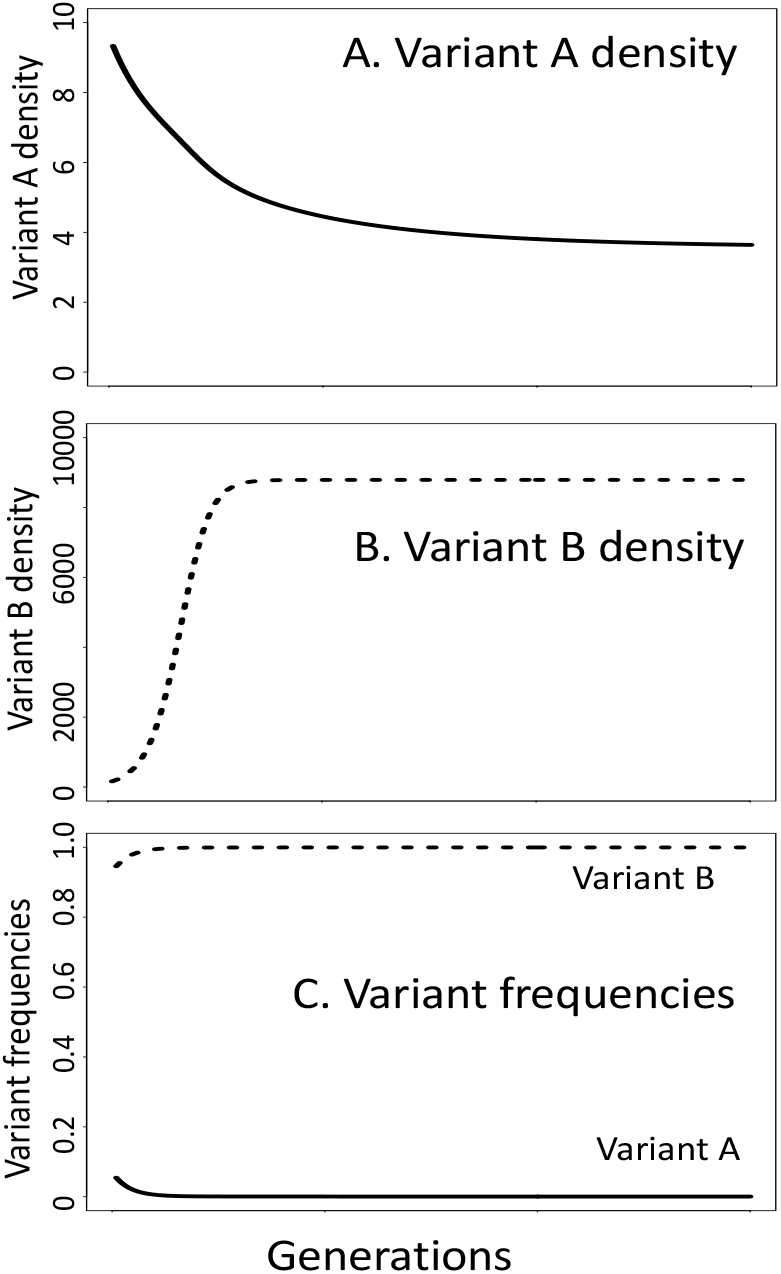
Multi-niche selection, an alternative model of balancing selection, is not a function of the core assumption of negative frequency-dependent selection models that relative fitness is a function of relative frequency in the population. Shown is a simulation where **variant A** has a selective advantage in **niche A** while **variant B** has a selective advantage in **niche B** (Supplemental material). Additionally, the carrying capacity in **niche A** is much lower than in **niche B** (K_A_ =10, K_B_=10000). At the start of the simulation, there are 10 **variant A** individuals (10% of the population) and 90 **variant B** individuals (90% of the population), yet the fitness (growth rate) of **variant A** individuals much lower than for **variant B** individuals. In the negative frequency-dependent selection model, the frequency of **variant A** should increase as it is currently less frequent than **variant B**. Although the conditions of negative frequency-dependent selection are not satisfied, both variants can be maintained in the population due to the selective advantage each enjoys in their preferred niche. Parameters used in the simulation: growth rate = 0.35, death rate in preferred niche = 0.05, death rate in non-preferred habitat =0.25, migration among niches = 0.01.

### Community diversity attributed to negative frequency-dependent selection

Prominent population geneticists including G.C. Williams and J. Maynard Smith, among many others, have demonstrated that the efficacy of natural selection decreases at increasing levels of biological organization such that selection among individuals within populations is much more efficient than selection among species within communities [91–93]. Additionally, selection at higher levels of organization (*i.e.* among species within communities) “tends to be undermined by natural selection at lower levels” (*i.e.* among individuals with populations) [94]. Nevertheless, several studies have suggested that negative frequency-dependent selection maintains species diversity within ecological communities. There is a rich empirical and theoretical history describing the causes and consequences of species diversity within ecological communities [2,95–99]. Mechanisms of coexistence function in two major ways: *equalizing* mechanisms minimize the average fitness differences between species while *stabilizing* mechanisms increase negative *intra*specific interactions relative to negative *inter*specific interactions [2]. Stabilizing mechanisms promote species coexistence and include mechanisms such as resource partitioning and frequency-dependent predation, as well as mechanisms that depend on spatial or temporal fluctuations in population densities or environmental factors. Equalizing mechanisms contribute to stable coexistence when they reduce large average fitness inequalities which might negate the effects of stabilizing mechanisms [2]. While some natural forces that affect the maintenance of community diversity have frequency-dependent mechanisms, this should not be mistaken for negative frequency-dependent selection which maintains polymorphisms within populations. Applying models of natural selection to levels of biological organization above the population level should be exercised only with the greatest caution [91].

The ‘Killing the Winner’ hypothesis is a recent endeavor to understand patterns of diversity within communities through a negative frequency-dependent selection framework [100,101]. More recent versions of the Killing the Winner hypothesis suggest that a frequency-dependent functional response in predator populations can promote community diversity. However, the functional response in this hypothesis is often not conditioned on the frequency of the species but on the presence or absence of character traits of the species that are being targeted by predators [101–104]. The “winner” in the Killing the Winner hypothesis refers to species that invest resource into reproduction at the expense of investing in predator defenses, which may or may not correspond to the most frequent species [102]. In these cases, neither the relative nor the absolute abundance of the prey species affects the functional responses of the predator.

## Conclusions

Understanding the processes that produce or maintain diversity in natural populations is a central challenge in evolutionary biology. Negative frequency-dependent selection maintains many noted and striking polymorphisms in nature [16,24,105–109], and many polymorphisms exist in the absence of a selective advantage favoring rare variants [30–35]. Ideally, one could unequivocally determine the causative process through observations of the patterns of variation in nature. Unfortunately, many processes result in identical patterns, especially when those patterns are observed over short time scales. In some cases, long-term observations of allelic dynamics can distinguish polymorphisms caused by mutation-selection balance or selection in a changing environment from a stable polymorphism resulting from balancing selection [33,110–114]. Evidence suggesting negative frequency-dependent selection - such as allelic cycles where each allele gains a selective advantage as it becomes more rare - may also be observed from long-term observational studies [24,115]. The patterns resulting from various evolutionary processes can also be tested through controlled and natural experiments such as manipulating allele frequencies in sub-populations [33,110,112,114].

Ecological and molecular mechanisms are rarely deducible from patterns [116], but accurate identification of the evolutionary processes causing the pattern can generate hypotheses about these mechanisms. For example, the northern acorn barnacle, *Semibalanus balanoides*, shows clear evidence of a balanced polymorphism at the mannose-6-phosphate isomerase (*mpi*) locus [117,118]. The pattern of *mpi* genotype frequencies among intertidal microhabitats, where one allele is common in high intertidal zones but rare in low intertidal zones, suggests that multi-niche selection maintains this polymorphism [119]. Experimental manipulations of genotypes among microhabitats confirmed that multiniche selection is the process responsible for the allelic variation [33,120]. The molecular mechanism linking mannose utilization with survivorship in high intertidal zones, where temperature and desiccation stress is high, was subsequently elucidated through controlled laboratory experiments [111]. As this and many other examples demonstrate, the ecological interaction or molecular mechanism underlying an evolutionary process can best be understood when the evolutionary process is accurately determined.

## Supplemental material

### Figure 1 data generated in R

~~~
### 3 genotypes S1S2, S1S3, S2S3
### S1 pollen can only pollinate S2S3 plant, 50% of time get S1S2, 50% S1S3
~~~

~~~
##Outline ###
## Start with Plants at different freq
## A pollen grain is chosen at random (by percent each is found in the population)
## pollen lands on 1 of the 100 plants (random number generator 1-100)
## Rejected if either allele of same plant is same as pollen
## if not rejected, makes seedlings with one of the 2 alleles (picked at random)
## repeat until 100 next generation plants
## repeat for 100 generations
~~~

~~~
##### PARAMETERS ####
S1S2<- 500 ## Starting pop of variant S1S2
S1S3<- 450 ## Starting pop of variant S1S3
S2S3 <- 50 ## Starting pop of variant S2S3
PopSize<-S1S2+S1S3+S2S3
S1<- (S1S2+S1S3)/(2*(S1S2+S1S3+S2S3)) ## starting number of S1 pollen grains
S2<- (S1S2+S2S3)/(2*(S1S2+S1S3+S2S3)) ## starting number of S2 pollen grains
S3<- (S2S3+S1S3)/(2*(S1S2+S1S3+S2S3)) ## starting number of S3 pollen grains
S1S2ng<-0
S1S3ng<-0
S2S3ng<-0
~~~

~~~
#### HOUSEKEEPING STUFF #####
generations<-5
pollens<-1000000
~~~

~~~
S1vector <- {}
S2vector <- {}
S3vector <- {}
~~~

~~~
S1vector[1] <- (S1S2+S1S3)/(2*(S1S2+S1S3+S2S3))
S2vector[1] <- (S1S2+S2S3)/(2*(S1S2+S1S3+S2S3))
S3vector[1] <- (S1S3+S2S3)/(2*(S1S2+S1S3+S2S3))
~~~

~~~
#### START OF MODEL SIMULATION #####
~~~

~~~
for (gens in 2:generations) { ## number of generations loop
pS1S2<-S1S2/(S1S2+S1S3+S2S3)
pS1S3<-S1S3/(S1S2+S1S3+S2S3)
pS2S3<-S2S3/(S1S2+S1S3+S2S3)
~~~

~~~
for (pol in 1:pollens) { ## mating loop
S1yes<-0
S2yes<-0
S3yes<-0
S1S2yes<-0
S2S3yes<-0
S1S3yes<-0
~~~

~~~
# choose pollen variant
~~~

~~~
pollenRand <-runif(1)
if (pollenRand<=S1) {S1yes<-1}
if (pollenRand > S1+S2) {S3yes<-1}
if (S1yes==0 & S3yes==0) {S2yes<-1}
~~~

~~~
# choose plant variant
PlantRand <-runif(1)
if (PlantRand<=pS1S2) {S1S2yes<-1}
if (PlantRand > pS1S2+pS1S3) {S2S3yes<-1}
if (S1S2yes==0 & S2S3yes==0) {S1S3yes<-1}
~~~

~~~
## next generation plants
if (S1yes == 1 & S2S3yes==1) {
      alleleSelect<-runif(1)
      if (alleleSelect<=.5){S1S2ng <- S1S2ng+1}
      else {S1S3ng <- S1S3ng+1}
}
~~~

~~~
if (S2yes == 1 & S1S3yes==1) {
      alleleSelect<-runif(1)
      if (alleleSelect<=.5){S1S2ng <- S1S2ng+1}
      else {S2S3ng <- S2S3ng+1}
^}^
if (S3yes == 1 & S1S2yes==1) {
      alleleSelect<-runif(1)
      if (alleleSelect<=.5){S1S3ng <- S1S3ng+1}
      else {S2S3ng <- S2S3ng+1}
}
~~~

~~~
#Stop when 100 seedlings
if (S1S2ng + S1S3ng + S2S3ng ==PopSize){break}
} ## mating loop
~~~

~~~
S1S2<-S1S2ng
S1S3<-S1S3ng
S2S3<-S2S3ng
S1S2ng <-0
S1S3ng <-0
S2S3ng <-0
~~~

~~~
S1<- (S1S2+S1S3)/(2*(S1S2+S1S3+S2S3))
S2<- (S1S2+S2S3)/(2*(S1S2+S1S3+S2S3))
S3<- (S2S3+S1S3)/(2*(S1S2+S1S3+S2S3))
~~~

~~~
S1vector[gens] <- (S1S2+S1S3)/(2*(S1S2+S1S3+S2S3))
S2vector[gens] <- (S1S2+S2S3)/(2*(S1S2+S1S3+S2S3))
S3vector[gens] <- (S1S3+S2S3)/(2*(S1S2+S1S3+S2S3))
~~~

~~~
}## end generations
~~~

~~~
S1S2
S1S3
S2S3
S1vector
S2vector
S3vector
~~~

~~~
minVect<-{}
minVect[1]<-min(S1vector)
minVect[2]<-min(S2vector)
minVect[3]<-min(S3vector)
~~~

~~~
maxVect<-{}
maxVect[1]<-max(S1vector)
maxVect[2]<-max(S3vector)
maxVect[3]<-max(S3vector)
~~~

~~~
minTotal<-min(minVect)
maxTotal<-max(maxVect)
~~~

~~~
minTotal
maxTotal
~~~

~~~
plot(S1vector,type = “l”, lwd=5, xlim=c(1, gens), ylim=c(minTotal-.01,maxTotal+.01), xaxt="n")
#axis(1, at = seq(1, 10, by = 1), las=2)
lines(S2vector,type = “l”, lty=3, lwd=5)
lines(S3vector,type = “l”, lty=4, lwd=5)
~~~

~~~
plot(S1vector,type = “l”, lwd=5, xlim=c(1, gens), ylim=c(minTotal-.01,maxTotal+.01), xaxt="n")
axis(1, at = seq(1, 10, by = 1), las=2)
plot(S2vector,type = “l”, lwd=5, xlim=c(1, gens), ylim=c(minTotal-.01,maxTotal+.01), xaxt="n")
axis(1, at = seq(1, 10, by = 1), las=2)
plot (S3vector,type = “l”, lwd=5, xlim=c(1, gens), ylim=c(minTotal-.01,maxTotal+.01), xaxt="n")
axis(1 at = seq(1, 10, by = 1), las=2)
~~~

~~~
#plot(S2vector,type = “l”, lty=3, lwd=5, xlim=c(1, gens), ylim=c(0,1))
#plot(dtvectorN1freq,type = “l”, lwd=5, xlim=c(0, gen), ylim=c(0,1))
~~~

~~~
#plot(dtvectorN1,type = “n”, xlim=c(1, gen), ylim=c(0,Ka))
#lo <- loess(dtvectorN1~time)
#xl <- seq(min(time),max(time), (max(time) - min(time))/1000)
#lines(xl, predict(lo,xl), col=‘black’, lwd=5)
~~~

~~~
##plot(dtvectorN2,type = “n”, xlim=c(1, gen), ylim=c(0,Kb))
#lo <- loess(dtvectorN2~time)
#xl <- seq(min(time),max(time), (max(time) - min(time))/1000)
#lines(xl, predict(lo,xl), col=‘black’, lwd=5)
~~~

~~~
#plot(dtvectorN1freq,type = “n”, xlim=c(0, gen), ylim=c(0,1))
#lo <- loess(dtvectorN1freq~time)
#xl <- seq(min(time),max(time), (max(time) - min(time))/1000)
#lines(xl, predict(lo,xl), col=‘black’, lwd=5)
~~~

~~~
#plot(dtvectorN2freq,type = “n”, xlim=c(0, gen), ylim=c(0,1))
#lo <- loess(dtvectorN2freq~time)
#xl <- seq(min(time),max(time), (max(time) - min(time))/1000)
#lines(xl, predict(lo,xl), col=‘black’, lwd=5)
~~~

## Appendix 1 Figure 3 data generated in R

~~~
### Make a model with 2 niches (A and B) and two variants (1 and 2)
### show how it is not frequency but abundance and carrying capacity that affect relative fitness
~~~

~~~
### cycle through differential fitness values (death rates in home vs away areas) and migration
###### values show when relative fitness changes and when polymorphism maintained
~~~

~~~
##### PARAMETERS ####
N1aStart<- 10 ## Starting pop of variant 1 (all start in their home niche)
N2bStart<- 100 ## Starting pop of variant 2 (all start in their home niche)
N1bStart <- 0 ## Starting pop of variant 1 in niche b
N2aStart <- 0 ## Starting pop of variant 2 in niche a
N1m <-0 ## migrant pool
N2m <-0 ## migrant pool
Ka<- 10 ## Carrying capacity of niche a
Kb<- 10000 ## Carrying capacity of niche b
rh <- .35 ### growth rate in correct niche
ra <-rh ## growth rate in incorrect niche (cycle through this in for loop)
## Birth rate same in both areas and controlled by K, death rate much higher in away (and happens
first)
dh<-.05
##da – cycles
generations<-150
~~~

~~~
#### HOUSEKEEPING STUFF #####
#i <- seq(.1, .5, by=.01)
#j <- seq(0, .3, by=.01)
k <- seq(0, generations, by=1)
i <- seq(.25, .25, by=.01)
j <- seq(.01, .01, by=.01)
~~~

~~~
dtvectorN1<-{}
dtvectorN2<-{}
dtvectorN1freq<-{}
dtvectorN2freq<-{}
time<-{}
~~~

~~~
#### START OF MODEL SIMULATION #####
~~~

~~~
for (da in i) { ##Selection differential loop

        for (m in j) { ## migration loop
~~~

~~~
N1a<- N1aStart ##resetting the starting population
N2b<- N2bStart
N1b<- N1bStart
N2a<- N2aStart
~~~

~~~
for (gen in k) {## generations loop
~~~

~~~
#Deaths
N1a <- N1a * (1-dh) ## deaths in the home area (where fitness is higher)
N1b <- N1b * (1-da) ## deaths in the away area (where fitness is lower)
~~~

~~~
N2a <- N2a * (1-da)
N2b <- N2b * (1-dh)
~~~

~~~
#Births
N1a <- N1a * (1+rh* (1 - (N1a+N2a)/Ka))
N1b <- N1b * (1+ra* (1 - (N1b+N2b)/Kb))
~~~

~~~
N2a <- N2a * (1+ra* (1 - (N1a+N2a)/Ka))
N2b <- N2b * (1+rh* (1 - (N1b+N2b)/Kb))
~~~

~~~
#migration
N1aT<- N1a - N1a*m ##emigrants leaving the population
N1bT<- N1b - N1b*m
N2aT<- N2a - N2a*m
N2bT<- N2b - N2b*m
~~~

~~~
if (N1aT < 0) {N1aT<-0}
if (N2aT < 0) {N2aT<-0}
if (N1bT < 0) {N1bT<-0}
if (N2bT < 0) {N2bT<-0}
~~~

~~~
N1am <- m * (N1a + N1b)/2 ## immigrants joining a population
N1bm <- N1am
N2am <- m * (N2a + N2b)/2
N2bm<-N2am
~~~

~~~
NaSpace<- Ka-N1aT-N2aT ## migrants cannot displace the residents
NbSpace<- Kb-N1bT-N2bT## migrants cannot displace the residents
if(NaSpace<0){NaSpace<-0}
if(NbSpace<0){NbSpace<-0}
~~~

~~~
if ((N1am+N2am) > NaSpace){
N1a <- N1aT + NaSpace*N1am/(N1am+N2am)
N2a <- N2aT + NaSpace*N2am/(N1am+N2am)
} else {
N1a <- N1aT + N1am
N2a <- N2aT + N2am
}
~~~

~~~
if ((N1bm+N2bm) > NbSpace) {
N1b <- N1bT + NbSpace*N1bm/(N1bm+N2bm) ## if the number of immigrants is too large, a
proportion join the population
N2b <- N2bT + NbSpace*N2bm/(N1bm+N2bm)
} else {
N1b <- N1bT + N1bm
N2b <- N2bT + N2bm
}
~~~

~~~
N1<-N1a+N1b
N2<-N2a+N2b
~~~

~~~
dtvectorN1[gen]<-N1
dtvectorN2[gen]<-N2
dtvectorN1freq[gen]<-N1/(N1+N2)
dtvectorN2freq[gen]<-N2/(N1+N2)
time[gen]<-gen

        } ## end generations loop
       } ##end migration loop
}## end selection loop
~~~

## Appendix 2 Ordinary differential equation (ODE) approximations of the simulation in Appendix 1

In the multiple niche selection model of balancing selection, the selective value of an allele is conditioned on its ability to exploit different environmental features in a heterogeneous habitat [59,60]. This simple ODE model is illustrative of a multiple niche polymorphisms that can maintain a stable polymorphism without selection favoring relatively rare variants.

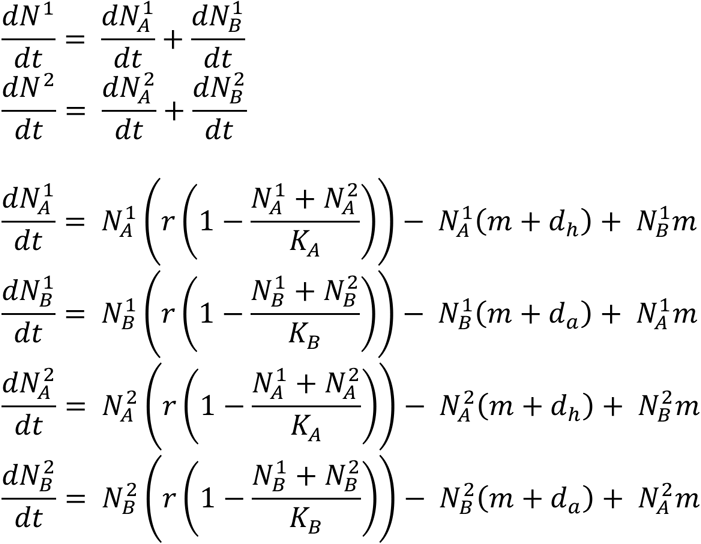

where *N*^1^ is the total abundance of variant 1, 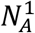 is the abundance of variant 1 in habitat *A* (its preferred habitat), *K_A_* is the carrying capacity of habitat *A, d_h_* and *d_a_* are the death rates in the preferred (*home*) and non-preferred (*away*) habitats, and *r* and *m*: are the intrinsic growth and migration rates (which do not differ among variants). Substituting parameter values similar to those used in the simulation (*r* = 0.35; *K_A_* = 100; *K_B_* = 1000; *d_h_* = 0.05; *d_a_* = 0.25; *m* = 0.01), a stable equilibrium can be found at *N*^1^ = 45 and *N*^2^ = 869 (*p* = 0.049; *q* = 0.951). The derivatives around *N*^1^ = 100; *N*^2^ = 500 (*p* = 0.167; *q* = 0.833) are negative for 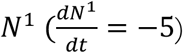 and strongly positive for 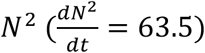, which is inconsistent with the expectation under negative frequency-dependent selection in which the rare variant should increase in frequency due to its rare advantage. This ODE model is qualitatively similar to one published in Ravigne *et al* (details in Appendix 1 of [60]):

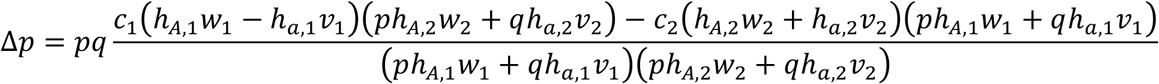

where *C*_1_ is the proportion of habitat 1, *w*_1_ is the viability of *A* in habitat 1 (where *A* has a survival advantage), *v*_1_ is the viability of a in habitat 1, and *h*_*A*,1_ is the habitat preference of *A* for habitat 1. Substituting parameter values similar to those used above (*c*_1_ = 0.99; *C*_2_ = 0.01; *h_A,1_ = h_a,2_* = 0.9999; *h*_*a*,1_ = *h*_*A*,2_ = 0.0001; *w*_1_ = *v*_2_ = 5; *w*_2_ = *v*_1_ = 1), a stable equilibrium can be found at *p* = 0.99 and *q* = 0.01. The derivative around *p* = 0.9 are nevertheless positive for *p* (*Δp* = 0.09), despite being at a much higher frequency, suggesting that the relative selective value of *p* does not become negative due to an increase in frequency.

